# Pontin/Reptin-associated complexes differentially impact plant development and viral pathology

**DOI:** 10.1101/2021.11.29.470430

**Authors:** Snigdha Chatterjee, Min Xu, Elena A. Shemyakina, Jacob O. Brunkard

## Abstract

Pontin and Reptin are essential eukaryotic AAA+ ATPases that work together in several multiprotein complexes, contributing to chromatin remodeling and TARGET OF RAPAMCYIN (TOR) kinase complex assembly, among other functions. Null alleles of *pontin* or *reptin* are gametophyte lethal in plants, which has hindered studies of their crucial roles in plant biology. Here, we used virus-induced gene silencing (VIGS) to interrogate the functions of *Pontin* and *Reptin* in plant growth and physiology, focusing on *Nicotiana benthamiana*, a model species for the agriculturally significant *Solanaceae* family. Silencing either *Pontin* or *Reptin* caused pleiotropic developmental and physiological reprogramming, including aberrant leaf shape, reduced apical growth, delayed flowering, increased branching, chlorosis, and decreased spread of the RNA viruses *Tobacco mosaic virus* (TMV) and *Potato virus X* (PVX). To dissect these pleiotropic phenotypes, we took a comparative approach and silenced expression of key genes that encode subunits of each of the major Pontin/Reptin-associated chromatin remodeling or TOR complexes (*INO80*, *SWR-C/PIE1*, *TIP60*, *TOR*, and *TELO2*). We found that many of the *pontin/reptin* phenotypes could be attributed specifically to disruption of one of these complexes, with *tip60* and *tor* knockdown plants each phenocopying a large subset of *pontin/reptin* phenotypes. We conclude that Pontin/Reptin complexes are crucial for proper plant development, physiology, and stress responses, highlighting the multifaceted roles these conserved enzymes have evolved in eukaryotic cells.

## Introduction

Pontin and Reptin are deeply conserved eukaryotic AAA+ ATPases that form a heteromeric complex with diverse biological roles (Huen *et al.*, 2010; Rosenbaum *et al.*, 2013; Dauden *et al.*, 2021). Clinical and mechanistic studies have implicated Pontin/Reptin in the metabolic and epigenetic regulation of various human diseases, including several cancers (Huber *et al.*, 2008; Flavin *et al.*, 2011; Osaki *et al.*, 2013; Mikesch *et al.*, 2018; Assimon *et al.*, 2019; Yan *et al.*, 2019; Armenteros-Monterroso *et al.*, 2019; Shin *et al.*, 2020; Lin *et al.*, 2020). Pontin/Reptin also contribute to human immune systems through roles in immune cell development and immunity signaling pathways (Arnold *et al.*, 2012; Hosokawa *et al.*, 2013; Zhang *et al.*, 2021). The importance of Pontin/Reptin to human health has prompted significant research effort to define their molecular functions, most of which can be broadly divided into two categories: (i) scaffolding chromatin remodeling complexes and (ii) co-chaperoning HSP90-mediated multiprotein complex assembly. In chromatin, Pontin/Reptin associate with the H2A/H2A.Z-substituting nucleosome remodeling complexes INO80 and SWR-1 (Kobor *et al.*, 2004), with the histone H4 acetyltransferase TIP60 complex (Ikura *et al.*, 2000; Jha *et al.*, 2008, 2013), and with telomerase (Venteicher *et al.*, 2008; Schořová *et al.*, 2019). Pontin/Reptin ATPase activity is apparently not required for many of their roles in chromatin remodeling (Yenerall *et al.*, 2020). In contrast, Pontin/Reptin complexes provide ATPase activity in their roles as co-chaperones for HSP90 in the R2TP (alias PAQosome) complex that promotes the assembly of functional snoRNPs (small nucleolar ribonucleoproteins) and PIKK (phosphatidylinositol kinase-like kinase, including TARGET OF RAPAMYCIN or TOR) complexes (Zhao *et al.*, 2008; Hořejší *et al.*, 2010; Boulon *et al.*, 2010; Kakihara & Houry, 2012; Kakihara *et al.*, 2014; Rivera-Calzada *et al.*, 2017; Martino *et al.*, 2018; Houry *et al.*, 2018; Maurizy *et al.*, 2018; Muñoz-Hernández *et al.*, 2019; Yenerall *et al.*, 2020; Coulombe *et al.*, 2020).

Evolutionarily, Pontin (aliases include RuvBL1, RVB1, and Tip49a) and Reptin (aliases include RuvBL2, RVB2, and Tip48/Tip49b) are distant homologues of the bacterial DNA-dependent ATPase RuvB, which participates in Holliday junction resolution. Within eukaryotes, Pontin and Reptin are remarkably conserved: to illustrate, Arabidopsis Pontin and Reptin are each ~90% similar and ~75% identical to their human orthologues. Much less is known about Pontin/Reptin in plants than in humans, however. Pontin was identified in yeast two-hybrid screens as a protein interactor of Arabidopsis NOD-like receptors (NLRs) RPM1 and RPP2, which confer resistance to virulent strains of *Pseudomonas syringae* (bacteria) and *Peronospora parasitica* (oomycetes), respectively. The possible role of Pontin and Reptin in NLR-mediated disease resistance was not mechanistically established, although mildly decreasing *Pontin* expression did perturb development and enhance disease resistance. Pontin and Reptin were also identified in Arabidopsis telomerase complexes and are apparently required for telomere maintenance (Schořová *et al.*, 2019).

Recently, we identified *Reptin* in a forward genetic screen for mutants with increased plasmodesmatal (PD) transport (Brunkard *et al.*, 2020). *pontin* and *reptin* knockouts are female gametophyte-lethal, but a weak allele of *reptin* (called *reptin-1* or *ise4*) carrying a missense mutation adjacent to the ATP-binding Walker A motif of the Reptin ATPase, A81V, can survive embryogenesis as a homozygote until the midtorpedo stage, when development arrests under standard growing conditions. Subsequent transcriptomic and functional investigation of *reptin-1* revealed that Reptin is required for TOR activity in plants, and that TOR is a crucial regulator of cell-cell trafficking through PD (Brunkard *et al.*, 2020). PD are nanoscopic membrane-bound channels in plant cell walls that connect the cytosol of neighboring cells, transporting metabolites, signaling molecules, small RNAs, proteins up to ~80 kDa, and viruses between neighboring cells (Brunkard & Zambryski, 2017; Azim & Burch-Smith, 2020). TOR is a broadly conserved eukaryotic protein kinase in the atypical PIKK family that coordinates eukaryotic metabolism in complex with its conserved interacting partners, RAPTOR and LST8 (Valvezan & Manning, 2019; Liu & Sabatini, 2020; Brunkard, 2020). TOR complex (TORC1) assembly and stability is dependent on interaction with the Pontin/Reptin co-chaperone complex, R2TP (Hořejší *et al.*, 2010; Kim *et al.*, 2013; Brunkard *et al.*, 2020; Yenerall *et al.*, 2020; Pal *et al.*, 2021), and both Pontin and Reptin have been confirmed as strong interactors of the TOR complex in plants (Van Leene *et al.*, 2019).

The early lethality of even weak loss-of-function alleles of *pontin* and *reptin* has limited our understanding of the significance of these genes at later developmental stages. Here, we use post-embryonic gene silencing to define the pleiotropic phenotypes impacted by loss of Pontin and Reptin. Then, we systematically silence critical subunits of the various established Pontin/Reptin-interacting complexes, including TOR, TOR-associated TELO2, and chromatin remodelers INO80, SWR1/PIE, and TIP60, and compare these phenotypes to the *pontin/reptin* knockdowns. We discover that Pontin and Reptin influence a broad range of plant phenotypes, including many unanticipated effects on morphology, physiology, flowering time, and pathogen defense, and, through comparison, hypothesize that most of these phenotypes can be explained by disruption of specific Pontin/Reptin-associated complexes. These findings substantially advance our understanding of the evolution of Pontin/Reptin biology in eukaryotes.

## Materials and Methods

### Plant Materials and Growth Conditions

*N. benthamiana* Nb-1 and *A. thaliana* Col-0 plants were grown under standard conditions with 16-h day/8-h night at ~120 μmol photons m^−2^ s^−1^ of photosynthetically active radiation. The reference Nb-1 genotype obtained from the Boyce Thompson Institute was used for all experiments (Bombarely *et al.*, 2012).

### Molecular Cloning and TRV-mediated VIGS

Virus-induced gene silencing (VIGS) vectors were prepared as previously described (Brunkard *et al.*, 2015; Horner & Brunkard, 2021), using oligonucleotides listed in Supplementary Table S1. RNA was isolated from Nb-1 shoots with the Spectrum Plant Total RNA (Sigma-Aldrich). cDNA was synthesized from RNA using random hexamers and SuperScript III reverse transcriptase (Fisher Scientific). Silencing triggers were amplified with Phusion DNA polymerase (New England Biolabs), digested alongside pYL156 with restriction enzymes as listed in Supplementary Table S1 (enzymes from New England Biolabs), and ligated with Promega T4 DNA ligase (Fisher Scientific). Ligations were transformed into house-made chemically competent XL1-Blue *Escherichia coli*. Kanamycin-resistant bacterial colonies were screened by colony PCR for positive clones. Plasmids were then miniprepped (Bioneer) and Sanger sequenced to confirm insert sequences. Manufacturer’s protocols were followed throughout. All constructs were then transformed into *Agrobacterium tumefaciens* GV3101 for plant transformation. Three week old plants were used for VIGS as previously described (Horner & Brunkard, 2021). The first true leaves were agroinfiltrated, with TRV::*GUS* as a negative (“mock”) control and TRV::*PDS* as a positive control to visually track silencing efficiency. Silencing was consistently observed within 14 days post-VIGS infiltration.

### Chlorophyll extraction

Chlorophylls were extracted as previously described (Müller-Moulé *et al.*, 2002). Briefly, three leaf punches per leaf were collected, flash frozen in liquid nitrogen, and then the frozen leaf punches were then ground with a plastic pestle. 100 μL of 100% acetone was then added, vortexed, and centrifuged at full speed for 1 min. This was step was done twice. 5μL of the resulting supernatant was taken and added to 995μL of cold, 80% acetone, vortexed, and centrifuged. Quantification was done using a spectrophotometer at 664nm for chlorophyll a and at 647 for chlorophyll b.

### Pathogen Inoculation

Four week old plants were used for all virus inoculations. One lower leaf per plant was syringe infiltrated with fresh overnight cultures of Agrobacterium resuspended in infiltration media (10 mM MgCl_2_, 10 mM MES, and 200 μM acetosyringone, pH 5.6) to a final OD_600nm_ = 0.5. Three Agrobacterium cultures were used that carried T-DNA encoding either TMV-GFP (Liu *et al.*, 2002; Burch-Smith *et al.*, 2012), PVX-GFP (Peart *et al.*, 2002), or a mock control. All plants were photographed seven days post agroinfiltration under a long wave UV lamp (100 watt, 365 nm, 115V-60Hz).

### Protein Extraction and Western Blot Assays

Leaves from VIGS plants infected with TMV-GFP were collected and snap frozen in liquid nitrogen. Protein was extracted from leaves in 100 mM MOPS (pH 7.6), 100 mM NaCl, 5% SDS, 0.5% β-mercaptoethanol, 10% glycerin, 2 mM PMSF, and 1× PhosSTOP phosphatase inhibitor (Sigma-Aldrich). Specific protein levels were assayed by Western blot protein using primary antibodies against GFP (Santa Cruz SC-9996) along with an HRP-conjugated goat anti-mouse IgG secondary antibody (Sigma Aldrich A4416).Total protein was visualized after transfer using Ponceau S red staining. All Western blot experiments were repeated at least three times, with representative results shown.

### RNA-seq

Tissue for RNA-seq was collected from TRV::*AtReptin,* TRV::*AtTIP60*, or mock-treated TRV::*GUS* plants 2 weeks post-infiltration with Agrobacterium cultures harboring the TRV binary vectors(Burch-Smith *et al.*, 2006). Plant leaves were collected and immediately frozen in liquid nitrogen. Total RNA was extracted using Trizol Reagent (Invitrogen, USA) from *N. benthamiana* leaves according to the manufacturer’s recommendations. Three pools of RNA from three individuals were prepared for the collections. RNA pools were made by combining 333.3 ng of RNA from each individual. Illumina TruSeq Stranded Total RNA kit with Ribo-Zero Plant was used to prepare libraries for RNA sequencing. Libraries were sequenced at the Vincent J. Coates Genomics Laboratory (QB3, Berkeley) using an Illumina Hi-Seq 4000 platform (150-bp paired-end run) and analyzed as previously described (Scarpin *et al.*, 2020). Reads were aligned with HISAT2 and counted with HTseq (Anders *et al.*, 2015; Kim *et al.*, 2015). Differential transcript abundance was determined with DESeq2 (Love *et al.*, 2014). Transcriptomes were further analyzed with Mapman software (Thimm *et al.*, 2004). Significantly affected gene categories were determined by MapMan using a Wilcoxon rank-sum test with the Benjamini-Hochberg-Yekutieli procedure to correct for the false discovery rate (Thimm *et al.*, 2004).

### Growth and development analyses of TRV-VIGS plants

Shoot phenotypes of plants infected with TRV carrying silencing triggers was performed as previously described (Busche *et al.*, 2021). Phenotypes were analyzed 2 weeks after agroinfiltrating to initiate VIGS, including counting branch number and quantifying chlorophyll levels. All pictures of leaf series were taken 2 weeks post VIGS. Flowering time data is reported for plants after plants flowered.

## Results

### Silencing *Reptin* or *Pontin* cause pleiotropic defects in shoots

We used a *Tobacco rattle virus* (TRV) VIGS system as a reverse genetics tool to knockdown *Pontin* and *Reptin* expression in *N. benthamiana* plants. Throughout, we used a mock silencing construct containing a fragment of the bacterial *GUS* gene (TRV::*GUS*) as a negative control and a silencing construct containing a fragment of the phytoene desaturase gene, *PDS* (TRV::*PDS*), as a positive control to confirm VIGS efficiency. Silencing *Pontin* or *Reptin* drastically decreased plant growth and caused pleiotropically aberrant leaf shape (Figure 1a & 1c). *Pontin* and *Reptin* knockdowns also displayed chlorotic leaves (Figure 1a). To quantify this phenotype, we extracted pigments from silenced leaves and measured total chlorophyll a and b spectrophotometrically. TRV::*Reptin* and TRV::*Pontin* plants accumulated significantly less amounts of both chlorophyll a and b, when compared to mock-treated plants (n = 9, p < 10^−3^, Student’s *t*-test) (Figure 1b).

**Figure 1.**
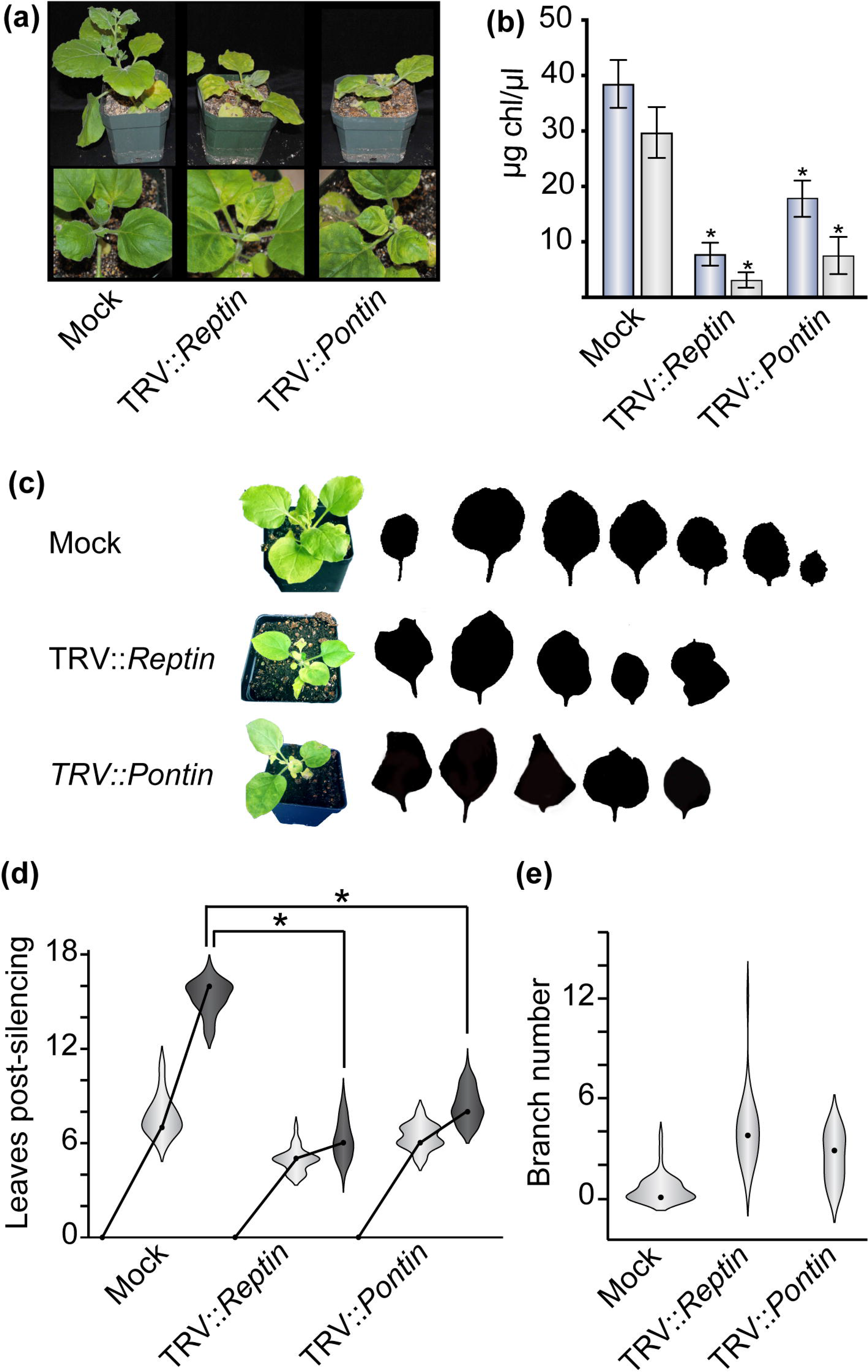
Silencing *Pontin* or *Reptin* causes comparable pleiotropic defects in shoot growth, development, and physiology. **(a)** Representative images of mock-treated (TRV::*GUS*) plants compared to TRV::*Reptin* and TRV::*Pontin* plants four weeks after initiating VIGS. **(b)** Knocking down *Reptin* or *Pontin* expression significantly decreases chlorophyll a and chlorophyll b content in leaves (*n* = 9, *p* < 10^−3^, Student’s *t*-test). Blue bars (left) indicate chlorophyll a levels, gray bars (right) indicate chlorophyll b levels. **(c)** TRV::*Reptin* and TRV::*Pontin* plants make fewer leaves with aberrant leaf shapes. Silhouettes are representative images taken from a leaf series of a single plant, with the first leaf that showed effects of silencing on the left and subsequent leaves ordered from oldest to youngest. **(d)** Leaf numbers were quantified 14 and 28 days after infiltrating with VIGS constructs. TRV::*Reptin* and TRV::*Pontin* plants made significantly fewer leaves than mock-treated (TRV::*GUS*) plants. **(e)** Lateral branches were counted for mock-treated (TRV::*GUS*), TRV::*Reptin*, and TRV::*Pontin* plants. Whereas most mock plants extended no or very few branches, TRV::*Reptin* and TRV::*Pontin* plants extended branches from a large number of axillary buds.

Since silencing *Pontin* and *Reptin* decreased growth in plants, we decided to count the number of emerged, expanding leaves starting soon after the onset of VIGS (two weeks after agroinfiltration) and again two weeks later to determine if these genes are required for leaf initiation and/or expansion. Mock-treated plants made 10.0 ± 0.2 leaves on average during the two weeks after agroinfiltration, whereas TRV::*Reptin* plants made only 5.0 ± 0.1 leaves and TRV::*Pontin* plants made only 6.7 ± 0.1 leaves, on average (n ≥ 51, p < 10^−3^, t-test). We went on to count leaves on the primary shoot every other day for the next 2 weeks. Whereas mock plants went on to make 15.2 ± 0.1 leaves on average 4 weeks post VIGS, TRV*::Reptin* and TRV*::Pontin* knockdown plants went on to make only 6.4 ± 0.2 and 8.5 ± 0.2 leaves, respectively (p < 10^−3^) (Figure 1d). The transition from vegetative to reproductive development was also delayed by silencing *Pontin* and *Reptin*. TRV::*Pontin* and TRV::*Reptin* plants flowered two or more weeks later than mock-treated plants, and many TRV::*Pontin* and TRV::*Reptin* knockdowns did not flower at all during our 4-week observation period. Despite reduced overall shoot growth, *Reptin-* and *Pontin-* silenced plants also made more branches when compared to mock-treated plants (Figure 1e).

We knocked down *Reptin* and *Pontin* in *Arabidopsis thaliana* and observed similar phenotypes. TRV::*AtReptin* and TRV::*AtPontin* plants were also smaller in size and displayed chlorosis, confirming that the effects of silencing *Reptin* and *Pontin* are not unique to *N. benthamiana* or its close asterid relatives. Importantly, in both species, TRV::*Pontin* plants were phenotypically indistinguishable from TRV::*Reptin* plants, suggesting that Reptin and Pontin’s impacts on plant development and physiology are largely (if not wholly) due to their roles as a heteromeric ATPase complex, rather than any independent functions as homomers.

### Silencing *TIP60* phenocopies many effects of knocking down *Pontin/Reptin*

To parse the phenotypes that we observed in TRV::*Reptin* and TRV::*Pontin* plants, we next used VIGS to knockdown expression of the genes that encode catalytic subunits of Pontin/Reptin-associated chromatin remodeling complexes, including *INO80*, *PIE1*, and *TIP60*. INO80 and PIE1 are involved in the replacement of H2A/H2A.Z/H2B dimers (Figure 2a), and mutants in *ino80* and *pie1* have been previously described in *A. thaliana* (Noh & Amasino, 2003; Fritsch *et al.*, 2004; March-Díaz *et al.*, 2008; Zhang *et al.*, 2015). TIP60 is a histone acetyltransferase (Figure 2a), also known as HAM1/HAM2 in *A. thaliana*; *TIP60* knockouts are female gametophyte lethal in *A. thaliana* (Latrasse *et al.*, 2008). Like *Reptin* and *Pontin* knockdowns, TRV::*NbTIP60* plants showed chlorotic leaves, which was confirmed by significantly lower quantities of chlorophyll a and chlorophyll b when compared to the mock-treated plants (n = 9, p = 10^−5^ and p = 0.003) (Figure 2c). TRV::*NbTIP60* plants also make less leaves compared to mock plants, about 6.0 ± 0.1 leaves on average, much like the Reptin and Pontin knockdowns (n ≥ 49) (Figure 2b & 2d).

**Figure 2.**
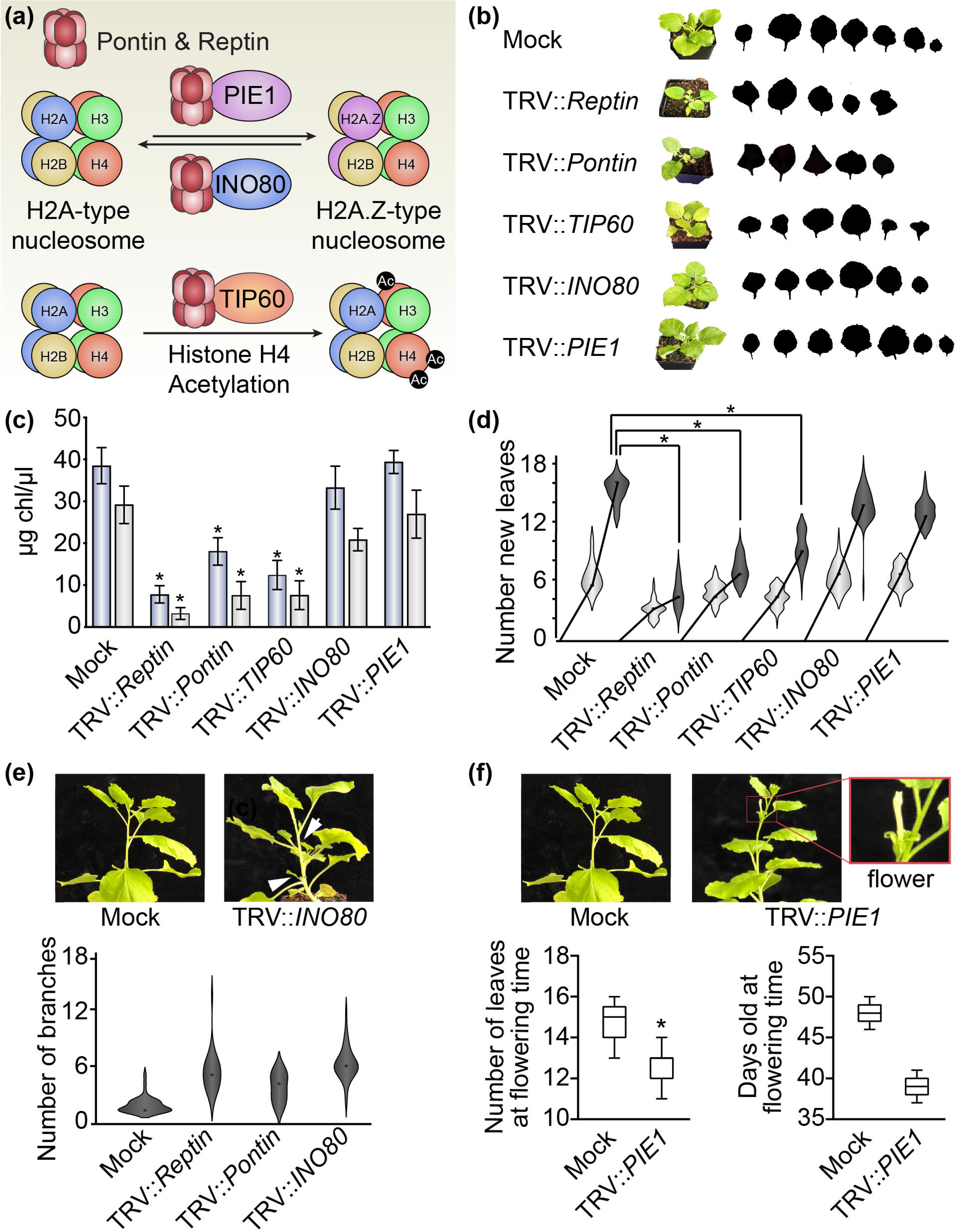
Silencing expression of Pontin/Reptin-associated chromatin remodeling complexes partially phenocopies TRV::*Reptin* and TRV::*Pontin* phenotypes. **(a)** Functional summary of the Pontin/Reptin-associated chromatin remodeling complexes targeted for silencing in this study. PIE1 and INO80 participate in exchanging H2A and H2A.Z subunits in nucleosomes, which can impact gene expression. TIP60 is an acetyltransferase that acetylates histone H4, which typically promotes gene transcription. **(b)** Representative eaf phenotypes of TRV::*TIP60*, TRV::*INO80*, and TRV::*PIE1* are shown as described in Figure 1a. TRV::*TIP60* somewhat disrupts leaf size and shape, whereas TRV::*INO80* and TRV::*PIE1* plants are largely similar to mock-treated (TRV::*GUS*) plants. **(c)** Leaf chlorophyll levels were measured for each silenced plant. Chlorophyll a (left, blue bars) and chlorophyll b (right, gray bars) concentrations are significantly reduced in TRV*::TIP60*, to similar levels as observed in TRV::*Reptin* and TRV::*Pontin* plants. TRV::*INO80* and TRV::*PIE1* plants showed no significant changes in chlorophyll levels. **(d)** Leaf numbers were tracked as described in Figure 1d. TRV::*TIP60* made significantly fewer leaves than mock-treated plants, similar to TRV::*Reptin* and TRV::*Pontin* plants. TRV::*INO80* and TRV::*PIE1* had no significant effect on leaf emergence rates. **(e)** TRV::*INO80* phenocopied the increased branching phenotype observed in TRV::*Reptin* and TRV::*Pontin* plants. Silencing *PIE1* and *TIP60* had no significant effect on branching in our experiments. **(f)** TRV::*PIE1* plants flowered significantly earlier than mock-treated (TRV::*GUS*) plants, similar to the early-flowering phenotype of *pie1* mutants in *A. thaliana*. In contrast, TRV::*Reptin* and TRV::*Pontin* plants rarely flowered during our observation period.

In contrast, neither TRV::*NbINO80* nor TRV::*NbPIE1* plants showed visible signs of chlorosis, and we did not detect any statistically significant change in chlorophyll a or b accumulation in the leaves of these plants (n= 9, p > 0.09) (Figure 2c). TRV::*NbINO80* and TRV::*NbPIE1* plants also did not show any difference in the number of leaves they developed compared to the mock plants (Figure 2b & d). Previous work suggested that *ino80* mutants display a mild increased branching phenotype in *A. thaliana* (Fritsch *et al.*, 2004). Similarly, TRV::*NbINO80* plants produced more branches (5.3 ± 0.3 branches) than TRV::*GUS* mock plants (0.6 ± 0.1 branches, *n* ≥ 51, *p* = 10^−8^), similar to TRV::*Reptin* and TRV::*Pontin* plants (Figure 2e). A screen for early flowering phenotypes identified *pie1* mutants in *A. thaliana* (Noh & Amasino, 2003), which led us to investigate TRV::*PIE1* flowering time. Indeed, knocking down *PIE1* in in *N. benthamiana* causes the plants to flower 9 ± 0.2 days earlier than mock plants (*n* ≥ 51, *p* < 10^−3^, Student’s *t*-test) (Figure 2f). This phenotype contrasts with the late flowering phenotype of TRV::*Reptin* and TRV::*Pontin* plants.

### Silencing *TOR* disrupts growth, similar to silencing *Pontin* and *Reptin*

Since Reptin and Pontin play a role in stabilizing the TORC1 dimer as integral members of the R2TP complex, we next used VIGS to knockdown *TOR* and another TOR-interacting co-chaperone, *TELO2,* to see if those genes could contribute to the pleiotropic phenotypes of *Reptin* and *Pontin* knockdowns. TELO2 is part of the TTT complex that contributes to the stabilization of TORC1 alongside R2TP (Figure 3a) (Takai *et al.*, 2007, 2010; Garcia *et al.*, 2017; Pal *et al.*, 2021). TRV::*NbTELO2* plants showed no apparent chlorosis and did not have significantly different accumulation of chlorophyll a or b compared to mock-treated plants (Figure 3c). Knocking down *TELO2* does not have strong impacts on plant growth and development, including leaf initiation rates, since the plants made 7.6 ± 0.1 leaves. (*n* ≥ 51, *p* = 0.1) (Figure 3b & 3d). Knocking down *TOR*, however, significantly decreased chlorophyll a and b accumulation by 34% for chlorophyll a and by 26% for chlorophyll b (*n* = 9, *p* < 0.002) (Figure 3c). TRV::*NbTOR* plants also showed severe dwarfism and made significantly fewer leaves compared to mock plants 2 weeks post VIGS (*n* ≥ 51, *p* < 10^−16^) and four weeks post VIGS (p < 10^−20^), much like the *Reptin* and *Pontin* knockdowns (Figure 3b & d). Therefore, disruption of TOR signaling could contribute to several of the pleiotropic phenotypes of TRV::*Reptin* and TRV::*Pontin* plants.

**Figure 3.**
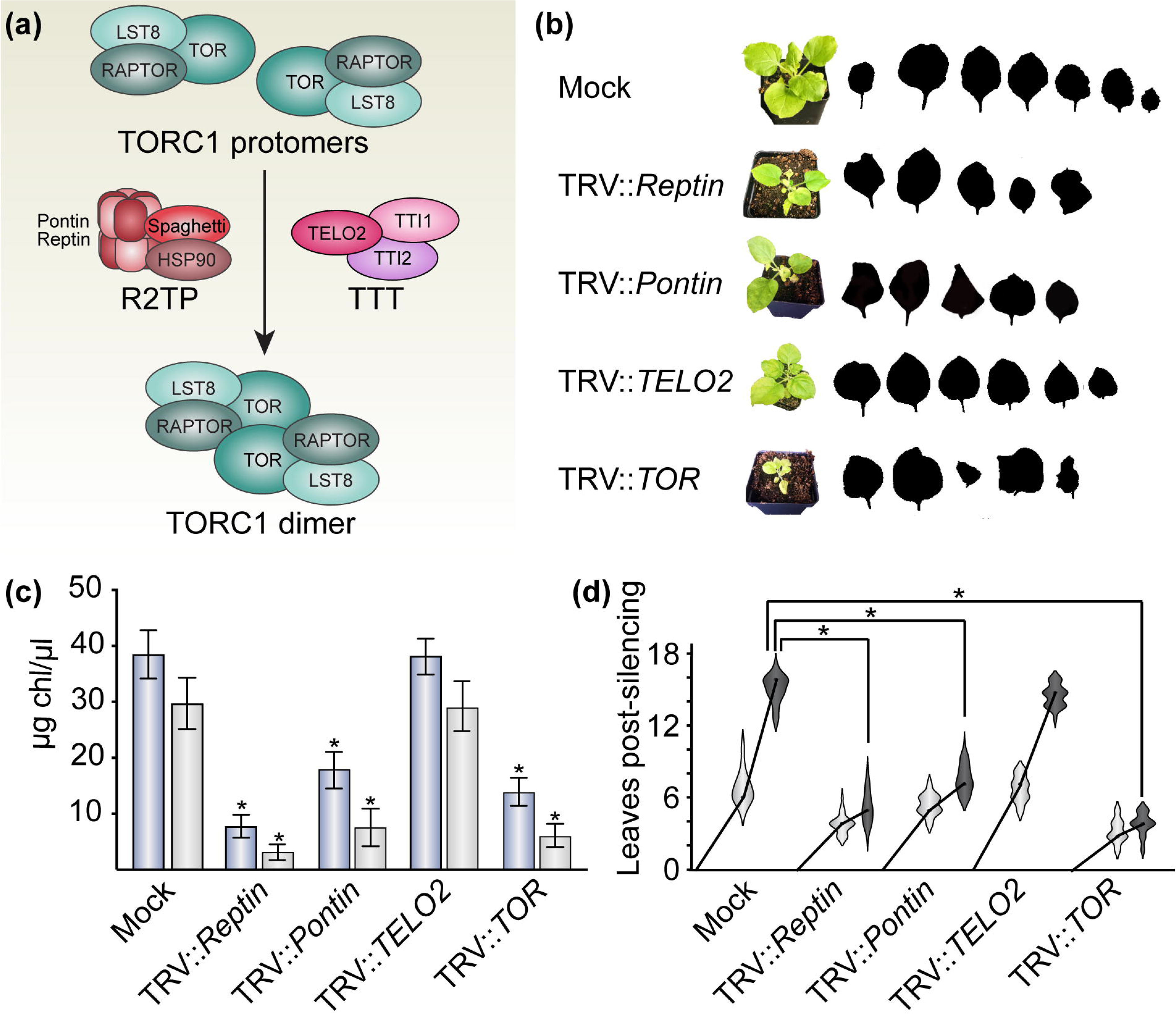
Silencing *TOR*, but not *TELO2*, severely disrupts plant growth and development with similar phenotypes to *pontin* and *reptin* knockdowns. **(a)** R2TP and TTT complexes are proposed to work in concert to assemble and/or stabilize the active TORC1 heterodimeric complex, as shown. **(b)** Shoot and leaf phenotypes for representative silenced plants, as described for Figure 1c. Silencing *TELO2* has minimal effects on leaf size and shape, whereas silencing *TOR* drastically reduces growth and causes aberrant leaf shapes, with phenotypes as severe as in TRV::*Reptin* and TRV::*Pontin* plants. **(c)** Chlorophyll levels are shown as in Figure 1b. Silencing *TOR* significantly reduces chlorophyll concentrations in leaves, but silencing *TELO2* has no significant effect. **(d)** Leaf initiation is not disrupted by silencing *TELO2*, but is almost completely arrested in TRV::*TOR* plants (panel as in Figure 1d).

### Silencing *Reptin, Pontin, TIP60*, or *TOR* restricts TMV infection

Pontin was first identified in plants as an interacting partner of *Arabidopsis thaliana* disease resistance proteins RPM1 and RPP5, which are both NOD-like nucleotide-binding leucine-rich-repeat receptors (NLRs), but differ in their N-terminal structure, which is either a coiled-coil domain (CC) or a Toll/Interleukin-1 Repeat-like domain (TIR), respectively (Whitham *et al.*, 1994; Grant *et al.*, 1995; Parker *et al.*, 1997; Holt *et al.*, 2002). The mechanistic relationship between Pontin/Reptin and the NLRs remains unclear, but *Pontin* knockdown plants exhibited increased resistance to the oomycete pathogen *Hyaloperonospora parasitica* (Holt *et al.*, 2002). Here, we tested whether knocking down *Pontin/Reptin* affects viral infection in plants. A *Tobacco mosaic virus* (TMV) strain expressing GFP was introduced into plants by agroinfiltration, and infection was tracked by UV-excited GFP fluorescence and by extracting protein from shoot apices and probing for viral GFP accumulation with α-GFP antibodies. By both methods, we observed that TMV movement was drastically restricted in TRV::*Reptin* and TRV::*Pontin* plants in contrast to mock plants (*n* ≥ 30) (Figure 4a and 4b). To determine whether this effect was specific to TMV or a more broad effect, we next infected plants with an unrelated RNA virus, *Potato virus X* (PVX), tagged with GFP. PVX-GFP infection was also strongly restricted in TRV::*Pontin* and TRV::*Reptin* plants (Supplementary Figure S1).

**Figure 4.**
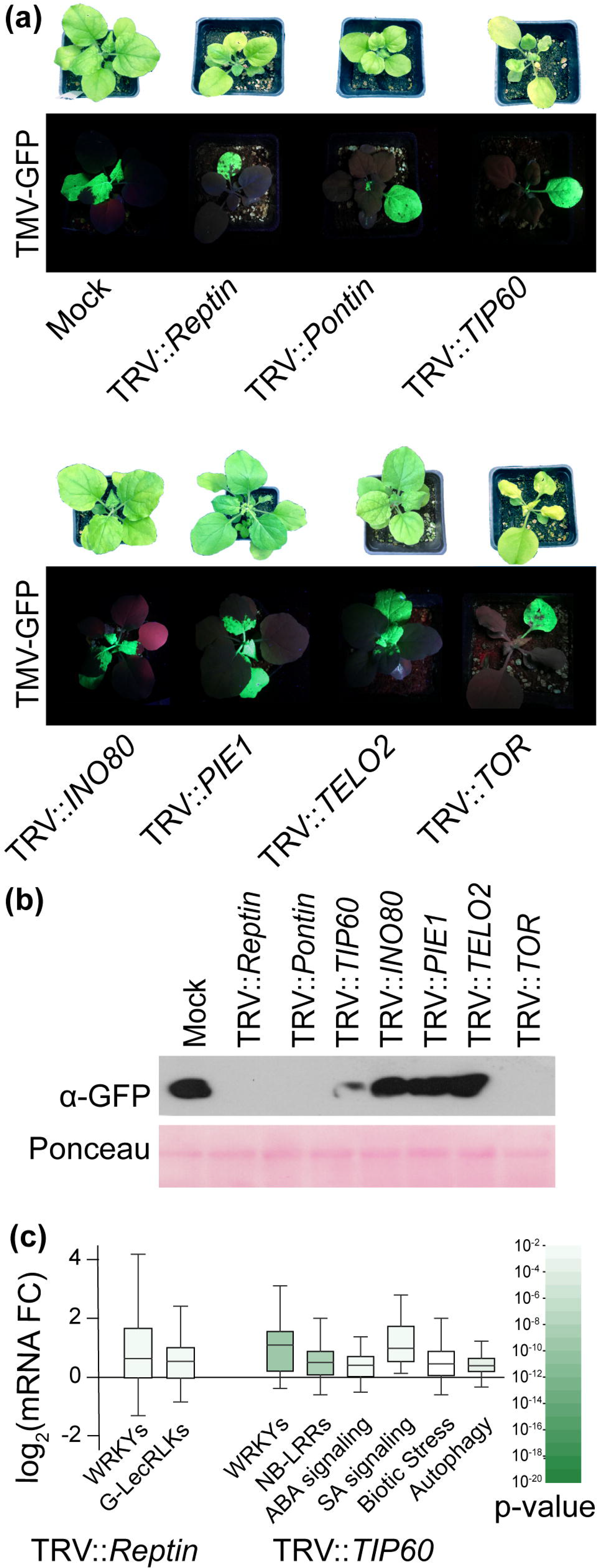
Pontin/Reptin-associated complexes impact viral pathogenesis and expression of biotic stress response pathways. **(a)** VIGS plants were infected with *Tobacco mosaic virus* or *Potato virus X* tagged with GFP (TMV-GFP or PVX-GFP, respectively) by agroinfiltration 14 days after initiating VIGS. Viral spread was observed using high-intensity UV light. Comparable results were obtained with either virus. Representative images of TMV-GFP spread are shown here 7 days after infection. TMV-GFP and PVX-GFP readily spread systemically in mock-treated (TRV::*GUS*), TRV::*INO80*, TRV::*PIE1*, and TRV::*TELO2* plants, but was restricted to the site of primary infection (lower leaf) in TRV::*Reptin*, TRV::*Pontin*, TRV::*TIP60*, and TRV::*TOR* plants. **(b)** Total protein was extracted from shoot apices of plants as in panel (a) and analyzed by SDS-PAGE followed Western blots against GFP. This validated the visual observation that TMV-GFP did not spread systemically in TRV::*Reptin*, TRV::*Pontin*, and TRV::*TOR*, and only slightly entered shoot apices in TRV::*TIP60* plants. Ponceau staining is shown as a loading control. **(c)** RNA-Seq of TRV::*Reptin* and TRV::*TIP60* plants showed consistent, significant upregulation of transcripts involved in biotic stress responses, including mRNAs that encode WRKY transcription factors, pathogen receptors, and stress response hormone signal transduction pathways. Gene ontologies were assigned with MapMan, *p* values were determined with MapMan using a Mann Whitney test with stringent false positive correction, and box plots indicate the quartiles and Tukey’s whiskers for all genes in each MapMan category.

Given the disrupted growth and enhanced resistance to viral infections of *pontin/reptin* knockdowns, we hypothesized that stress responses might be generally upregulated in these plants. Indeed, using RNA-Seq, we observed significant and broad induction of biotic stress/immune signaling response genes in *reptin* knockdown plants, which suggests that constitutive activation of the plant immune system might be responsible for the restriction of TMV and PVX spread in TRV:*Reptin* plants (Figure 4c, Supplementary Tables S2, S3). Incidentally, we had not observed any significant induction of biotic stress response genes in the *reptin-1* mid-torpedo stage embryo transcriptome (Brunkard *et al.*, 2020), which could be due to the different developmental stage or could suggest that this weak, partial loss-of-function allele of *reptin* has distinct effects on plant physiology from complete silencing of the *Reptin* gene.

We next tested whether the Pontin/Reptin-associated chromatin remodeling complexes might influence viral infections. We found that knocking down *INO80* or *PIE1* had no discernible effect on TMV or PVX infection, but that silencing *TIP60* delayed viral spread similarly to the restricted infections in TRV::*Pontin* and TRV::*Reptin* plants (Figure 4a). Since TMV and PVX are RNA viruses that replicate in the cytoplasm, and TIP60 is a chromatin-associated complex, the effects of TIP60 histone acetyltransferase activity on viral spread are most likely indirect due to changes in nuclear gene expression. Indeed, knocking down *TIP60* induced widespread expression of biotic stress response genes that could contribute to the restriction of TMV and PVX infection, correlating with the induction of biotic stress response genes in *Reptin* knockdown plants (Figure 4c).

Finally, we tested whether loss of TOR and/or TELO2 activity could contribute to the restricted viral spread in TRV::*Reptin* and TRV::*Pontin* plants. TOR activity is required for the replication and spread of many human pathogenic viruses (including coronaviruses (Zhou *et al.*, 2020; Karam *et al.*, 2021; Mullen *et al.*, 2021), herpesviruses (Chuluunbaatar *et al.*, 2010; Moorman & Shenk, 2010), and many more) and has been implicated in the replication of another family of plant RNA viruses, the potyviruses (Ouibrahim *et al.*, 2015). As with the other developmental and physiological phenotypes, knocking down *TELO2* had no discernable effect on TMV or PVX infection (Figure 4a). Silencing *TOR*, however, strongly prevented TMV and PVX spread, much like silencing *Reptin*, *Pontin*, and *TIP60* (Figure 4a). This experiment does not distinguish between direct effects of TOR signaling on TMV or PVX replication and spread versus indirect effects due to induction of biotic stress responses, although we should note that our previous work demonstrated that inhibiting TOR induces biotic stress responses in Arabidopsis (Scarpin *et al.*, 2020). Thus, we conclude that silencing TOR broadly restricts plant RNA viral spread, most likely by disrupting growth and broadly inducing biotic stress responses.

## Discussion

In this work, we interrogated the roles of the essential AAA+ ATPases Reptin and Pontin in post-embryonic plant development and physiology using a genetic knockdown approach, VIGS. We and others have previously used VIGS to investigate the role of essential genes that are required to survive embryogenesis and/or gametophyte development (Teresa Ruiz *et al.*, 1998; Stonebloom *et al.*, 2009; Burch-Smith & Zambryski, 2010; Ahn *et al.*, 2011; Brunkard *et al.*, 2020). Whereas silencing many embryo-lethal genes, including the regulators of plasmodesmatal transport *ISE1*, *ISE2*, and *ISE3*, causes relatively mild phenotypes (e.g., partial chlorosis, slightly reduced growth), silencing *Reptin* (alias *ISE4*) or *Pontin* causes severe, pleiotropic defects. This finding reflects the essential roles these genes play throughout the plant life cycle. To parse these pleiotropic phenotypes, we took a reverse genetic approach by silencing key components of several established Pontin/Reptin-associated complexes, namely the nuclear chromatin remodelers INO80, PIE1/SWR1, and TIP60 and the cytosolic metabolism-regulating complexes R2TP, TTT, and TORC1. Broadly, we found that most of the *pontin* and *reptin* knockdown phenotypes could be phenocopied by silencing one of these interactors.

The most severe phenotypes, including delayed leaf initiation rates, chlorosis, and severely misshapen leaf development, were consistently observed in *pontin*, *reptin*, *tip60*, and *tor* knockdowns. Unexpectedly, we did not observe similarly severe phenotypes in *telo2* knockdowns, although *telo2* is also essential for plant embryogenesis (Garcia *et al.*, 2017). Indeed, loss of many apparently important interacting partners of TOR, including the R2TP subunit Spaghetti (Brunkard *et al.*, 2020), the TORC1 subunit LST8 (Moreau *et al.*, 2012), and (as we show here) the TTT subunit TELO2, do not disrupt growth and development anywhere near as severely as disrupting TOR itself. Even in well-studied biomedical models, including yeast and human cell lines, some of these interactors are only conditionally required to maintain or support TORC1 activity; for example, mLST8, which is consistently found in the mTORC1 complex in wild-type cells, is apparently partially or wholly dispensable for mTORC1 activity under most circumstances (Guertin *et al.*, 2006; Hwang *et al.*, 2019). Multifaceted functional genetics approaches, including analysis of heritable mutations, acute gene silencing (e.g., VIGS), and chemical genetics, will be useful to determine how these genes contribute to metabolic regulation in response to environmental cues.

Strikingly, *pontin* and *reptin* knockdowns are phenotypically indistinguishable, which supports the prevailing hypothesis that Pontin and Reptin exclusively or, at least, most crucially function in a heteromeric ATPase complex. One of these genes cannot compensate for loss of the other, despite their remarkable similarity to each other and shared evolutionary origins as paralogous orthologues of bacterial RuvB. Although we were able to copy some of the *pontin* and *reptin* knockdown phenotypes by silencing their established interacting partners, none of these other knockdowns fully phenocopied the severity of *pontin/reptin*. One hypothesis to explain this result would be that the concerted loss of all of Pontin and/or Reptin’s functions is synergistically required to match the *pontin/reptin* knockdown phenotypes. Alongside this hypothesis, it remains likely that the full range of Pontin/Reptin functions in plant cells is not yet established. Ongoing efforts to directly define the molecular functions of these proteins directly in plants, rather than by orthology to conserved eukaryotic functions, could illuminate new or repurposed roles for the Pontin/Reptin ATPases in plants and explain some of the remaining pleiotropic phenotypes we observed.

Pontin/Reptin were first studied in plants following their discovery in a yeast two-hybrid screen for interactors of nucleotide-binding leucine-rich repeat (NB-LRR) NOD-like receptors (NLRs) that confer disease resistance in plants (Holt *et al.*, 2002). Although NLRs are typically activated in response to a limited range of specific pathogens (often only a subset of strains within a species), this early report found that slightly reducing *Pontin/Reptin* expression conferred resistance to at least some oomycete and bacterial pathogens. Here, we found that *pontin* and *reptin* knockdowns are much more resistant to viral pathogens as well, which we demonstrated with two unrelated RNA viruses, TMV and PVX. Moreover, transcriptomic analysis of *reptin* knockdowns confirmed that biotic stress responses are broadly induced in these plants, which would support the hypothesis that the Pontin/Reptin complex is required to limit spurious activation of the plant immune system. We also found that biotic stress responses are constitutively induced in *tip60* knockdowns, and *tip60* plants are accordingly also resistant to TMV and PVX. As with several other phenotypes, however, *tor* knockdowns shared similar phenotypes to *tip60*, *pontin*, and *reptin*, limiting spread of TMV and PVX. We recently showed that reducing TOR activity activates biotic stress responses, so this effect is also unlikely to be specific to just these two viruses, but instead an indication that silencing *tor* confers resistance to diverse pathogens. Indeed, another report using RNA interference to partially suppress *TOR* expression rice showed that *tor* knockdowns are also more resistant to the bacterial pathogen *Xanthomonas oryzae* (De Vleesschauwer *et al.*, 2018) Therefore, while we cannot propose a single mechanism by which knocking down *pontin* and/or *reptin* confers resistance to oomycetes, bacteria, and viruses, we do argue that these effects can be explained by disruption of either TIP60 or TOR activities, and are therefore not dependent of the previously observed protein-protein interactions between Pontin/Reptin and NLRs *in vitro* and in yeast.

In conclusion, this report demonstrates that *Pontin* and *Reptin* are crucial, multifunctional genes that impact a range of phenotypes in plants. Given the growing interest in Pontin and Reptin for their roles in human health, a deeper understanding of the evolutionary conservation of roles for Pontin/Reptin in cells, as well as the possible exaptation of new functions for these proteins in distinct lineages, could illuminate new lines of inquiry for the biomedical research community. More immediately, our results demonstrate that VIGS can be used to determine roles for gametophyte-lethal genes in postembryonic development and that comparative impacts of VIGS on multiple essential genes can be used to begin parsing how pleiotropic phenotypes are caused by disruption of promiscuous protein complexes.

## Supplementary data

Figure S1. Plants infected with PVX-GFP, shown as for TMV-GFP infections in Figure 4a.

Table S1. Oligonucleotide primers used in this study.

Table S2. RNA-Seq results for TRV::*Reptin* and TRV::*TIP60*.

Table S3. Significant MapMan gene ontologies of TRV::*Reptin* and TRV::*TIP60* RNA-Seq.

## Author Contributions

SC, MX, EMS, and JOB conceived the project, designed the experiments, performed the experiments, and analyzed data. SC and JOB wrote the manuscript.

## Acknowledgments

We thank Patricia Zambryski, M. Regina Scarpin, Michael Busche, Cynthia Amstutz, Samuel Leiboff, Tessa Burch-Smith, Barbara Baker, and Jeffrey Tung for reagents and support with conducting and designing experiments.

## Data availability

RNA-Seq sequencing data were deposited with NCBI, bioproject ID PRJNA736936.

